# Genomic study of oral lichen planus and oral microbiome with RNAseq

**DOI:** 10.1101/2020.02.12.946863

**Authors:** Evelyn F. Zhong, Andrea Chang, Andres Stucky, Xuelian Chen, Tarun Mundluru, Mohammad Khalifeh, Parish P. Sedghizadeh

## Abstract

Oral lichen planus (OLP) is a common chronic inflammatory disease affecting the oral mucosa. The pathogenesis of OLP is incompletely understood but is thought to be related to the immune system. As the oral cavity is a major reservoir and transmission gateway for bacteria, viruses, and fungi, the microbial composition of the oral cavity could play a role in the pathogenesis of OLP. However, due to limitations of analytic technology and incomplete knowledge of the microbial community in the oral cavity, it is not yet clear which pathogens are associated with OLP. Next-generation sequencing (NGS) is a powerful tool that can help to identify pathogens for many infectious diseases. In this study, we compared host cell gene expression profiles and microbial profiles from OLP patients and matched healthy individuals. We identified activation of the hepatocyte nuclear factor alpha (HNF4A) network in OLP patients and potential pathogens, including *Corynebacterium matruchotii, Fusobacterium periodonticum, Streptococcus intermedius, Streptococcus oralis*, and *Prevotella denticola. P. denticola* is capable of activating the HNF4A gene network. Our findings shed light on the previously elusive association of OLP with various diseases like hepatitis, and indicate that OLP is a T-helper type 17 (Th17)-mediated mucosal inflammatory process. The molecular pathways and microbes identified here can inform future investigations into OLP pathogenesis and development of novel therapeutics for OLP treatment.

## INTRODUCTION

Lichen planus is a common chronic mucocutaneous inflammatory disease affecting the skin, nails, eyes, urogenital tract, and oral mucosa. (1) Oral lichen planus (OLP) occurs specifically in the oral cavity. It is believed to affect the oral mucosa through T-cell mediated chronic inflammation, and some investigators have suggested that Th2-mediated inflammation can also contribute to the pathogenesis of OLP. (2)

Five clinical subtypes of OLP are usually seen: reticular, plaque-like, atrophic, erosive-ulcerative, and bullous. Symptomatic lesions can appear and regress over time. Cutaneous involvement in addition to oral mucosal lesions is seen clinically in a subset of patients. (3) The most commonly affected oral location affected, regardless of subtype, is the buccal mucosa, usually with symmetrical involvement.(4) This is followed by the gingiva, tongue, lip, and palate. OLP has a greater prevalence in females compared to males, and in females typically takes a more chronic course and has higher potential for significant morbidity. (5) OLP has a prevalence of approximately 0.5-2% with an age of onset between 30 and 60 years. (3)

A major clinical challenge for clinicians is providing patients with successful management or treatment of OLP without negative side effects. Currently, one of the most common treatments for OLP is topical corticosteroids. (6) Topical therapy is commonly favored over systemic therapy as the former is easier and more cost-effective than the latter. Systemic therapy often requires additional treatment such as concomitant topical therapy. (7) However, some patients do not respond to topical therapy. Also, the use of topical treatments without proper monitoring and evaluation can lead to oral candidiasis with associated burning mouth and hypogeusia, hypersensitivity reactions to the drug, inhibition of the hypothalamic-pituitary-adrenal axis and secondary adrenal insufficiency, or blood glucose dysregulation. Reports of patients with reactions to topical corticosteroids have also been detailed and highlight the challenges in managing patients with OLP. (8).

Investigating the gene networks of host cells and the microbiome associated with OLP could have a significant impact on our understanding and ultimately clinical treatment of OLP. In this study, we performed RNAseq to compare the host gene expression and microbial profiles of OLP patients with those of healthy controls. We identified specific host gene expression profiles of OLP as well as unique microbial profiles in OLP patients. These findings provide insights into OLP pathogenesis and could help inform future targeted therapies.

## METHODS

### Patient population

Institutional Review Board approval was obtained for this study (USC IRB # HS-16-00518). Inclusion criteria for OLP patients included males or females over age 50 with biopsy and histologically confirmed lichen planus diagnosis by a board-certified oral pathologist, and the ability to provide informed consent. Exclusion criteria for OLP patients included xerostomia or significant polypharmacy where saliva collection could be difficult or prohibitive, active systemic infection or oral infection such as periodontitis or abscess, detectable subgingival or supragingival plaque, active antibiotic therapy, and any history of skin involvement with lichen planus. Ascertainment criteria for controls included healthy male or female patients over age 50 without a diagnosis of cutaneous or oral lichen planus in any form, one or no comorbidities, no xerostomia or polypharmacy, no active systemic or oral/periodontal infection, and no detectable subgingival or supragingival plaque.

### Sample collection

Saliva and buccal mucosa wash samples of patients diagnosed with OLP (n=4) and healthy controls (n=4) were collected in OMNIgene saliva collection kits (Genotek). Saliva and buccal wash cells were collected from each patient under standardized conditions. Salivary flow rates vary significantly among individuals and in the same individual under different conditions or during different times of the day. Thus patients were instructed not to eat, clean, or rinse their mouth 1 hour before saliva collection, because these activities can affect the microbial environment. Saliva for all study subjects was collected at the same time of day using a draining method with the patient in an upright head position. Whole unstimulated saliva was collected for 5 minutes in a Proflow sialometer (Amityville, NY) or until 0.5 mL of saliva was collected (0.1 mL/min average). Samples were subsequently used for molecular studies.

### RNAseq

Libraries were constructed using Nextera DNA Flex Library Prep Kit (Illumina) and the Ribo-Zero rRNA Removal Kit (NEB). DNA and RNA were sequenced and results were analyzed using Partek Flow. To investigate the relationship between OLP and oral microbial status, we conducted DNA and RNA sequencing of patient saliva with buccal epithelial cells and aligned it to both human and bacterial viral genomes. We performed an observational (clinical paired with genomic patient information) study to identify possible associations. For differential expression analysis, we used Partek’s Gene Specific Analysis method. To identify significantly differentially expressed genes among different tissues from the same patient, a cutoff of FDR adjusted two-fold change was applied.

### Microbial profiling

Centrifuge is a taxonomic profiling tool (9) and was used to allocate unmapped reads to the microbial genomes and calculate the abundance of known microbial organisms for all samples. We used the -q flag to indicate that the input was in fastq format and all other arguments were set to their default values. All the available reference genomes in NCBI for bacterial and viral sequences were used for classification (up to January 4, 2018).

### PCR confirmation

PCR assays were used to confirm microbes identified from RNAseq. Primer for detection of *Prevotella denticola* was:

Forward Primers: TAATACCGAATGTGCTCATTTACAT

Reversed primer: TCAAAGAAGCATTCCCTCTTCTTCTTA

with an amplicon size of 316 bp (10).

## RESULTS

### Clinical and pathologic features

We studied 10 patients with OLP and five control patients without OLP; our OLP patient population had signs of both reticular and erosive patterns, and the most common sites of involvement were the buccal mucosa, gingiva, and tongue **(Table 1)**. A common clinical sign of OLP is oral lesions with radiating whitish gray lines or thread-like papules sometimes known as Wickham’s striae, which can be lacy or reticular, annular, patches, or strings. Desquamative gingivitis is another common finding with OLP **(Fig. 1 A&B)**. When Wickham’s striae predominate clinically, the term ‘reticular OLP’ is used. When atrophic or ulcerative lesions predominate clinically, the term ‘erosive OLP’ is used. Significant clinical characteristics of OLP often include lesions that alternate between periods of exacerbation and quiescence, as was seen in our patient population. (11) Patients may notice an irritation to the oral cavity and later develop a burning sensation. The burning sensation can be further aggravated by the intake of spicy or hot foods and can interfere with patient eating habits and quality of life. (12) None of our OLP patients had skin involvement or other tissue involvement outside the oral cavity.

**Table 1.** Clinicopathologic and demographic features of the study population cases and controls.

| **Cases** | **Age** | **Sex** | **Clinical oral diagnosis** | **Comorbidities** | **OLP Treatment** |
| --- | --- | --- | --- | --- | --- |
| **1** | 61 | F | OLP (reticular) buccal mucosa bilateral and gingiva generalized | None | Dexamethasone oral elixir 0.5mg/mL rinse and spit tid |
| **2** | 76 | M | OLP (erosive) buccal mucosa bilateral and gingiva generalized | Hypertension | Clobetasol gel 0.05% for topical use tid |
| **3** | 62 | F | OLP (reticular) buccal mucosa bilateral and gingival generalized | Hearing impairment | Fluocinonide gel 0.05% for topical use tid |
| **4** | 63 | F | OLP (erosive and reticular) buccal mucosa bilateral and gingiva generalized | Hypertension, angina | Fluocinonide gel 0.05% for topical use tid & dexamethasone as in case 1 |
| **5** | 61 | F | OLP (reticular) buccal mucosa bilateral | Mitral valve prolapse, anemia | Dexamethasone as in case 1 |
| **6** | 63 | F | OLP (erosive and reticular) buccal mucosa bilateral and tongue | Hypothyroidism, depression | Dexamethasone as in case 1 |
| **7** | 59 | F | OLP (erosive and reticular) buccal mucosa bilateral and gingiva generalized | Anxiety | Dexamethasone as in case 1 |
| **8** | 72 | F | OLP (erosive and reticular) buccal mucosa bilateral | Fibromyalgia, osteoarthritis, hypertension, hypothyroidism | Dexamethasone as in case 1 |
| **9** | 78 | F | OLP (reticular) buccal mucosa bilateral and tongue | Hypertension, DMII | None |
| **10** | 79 | F | OLP (reticular) buccal mucosa bilateral | Hypothyroidism, osteoporosis | Dexamethasone as in case 1 |
| **Controls** |  |  |  |  |  |
| **1** | 51 | F | Simple bone cyst | None | N/A |
| **2** | 69 | F | Fractured dental filling | Hypertension | N/A |
| **3** | 72 | F | Psychogenic bruxism | None | N/A |
| **4** | 59 | M | Dental crown fracture | Hypertension | N/A |
| **5** | 57 | M | None | None | N/A |

**Fig 1.**
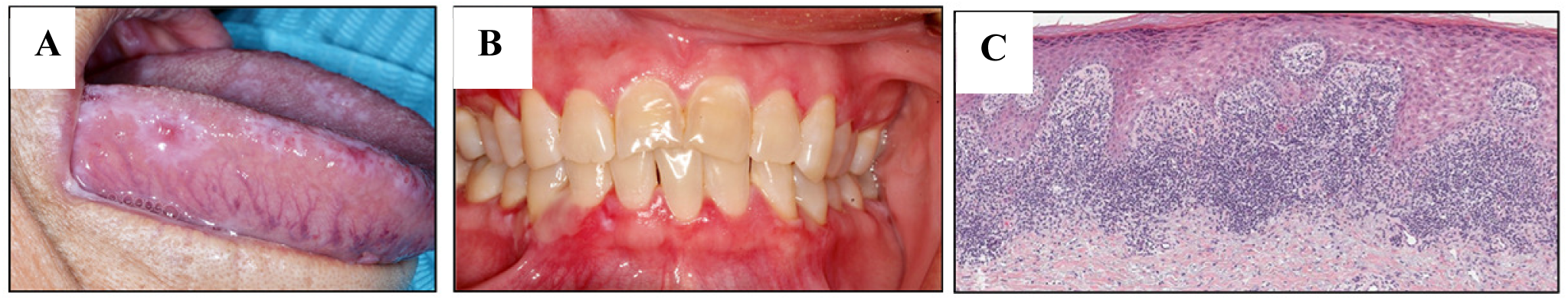
Clinical and pathologic features of OLP. A) image shows OLP involving the tongue with characteristic white striae (reticular variant) and red atrophic lesions (erosive variant). B) image shows OLP with inflamed and desquamative gingivitis. C) Histopathology of lichen planus demonstrates the characteristic band-like inflammatory cell infiltrate subjacent to the epithelium with “saw-tooth” rete ridge morphology and hydropic degeneration of basal keratinocytes.

It is usually recommended to confirm the diagnosis of OLP with an oral biopsy and histopathological examination, as lesions may mimic dysplasia or malignancy in addition to other conditions. (13) Thus, we confirmed OLP diagnosis in all studied patients via histopathologic evaluation. Histopathologic criteria for diagnosis include the presence of a well-defined, band-like zone of inflammatory cell or lymphocytic infiltration within the superficial part of the connective tissue or lamina propria **(Fig. 1C)**. Histologic evidence of hydropic degeneration in the basal cell layer with absence of epithelial dysplasia is also common. (14) The finding of “saw-tooth”-shaped rete ridges and colloid bodies are other useful features which help support the diagnosis of OLP.

### Molecular analysis of OLP-related gene network

The pathology of OLP is strongly related to immune dysregulation, like any other autoimmune disease. The most popular theory is that activated cytotoxic CD8+ cells target basal keratinocytes, after which CD4+ helper T cells secrete TH1 cytokines. (15) Along with T cells, growth factors, inflammatory and proapoptotic mediators mediate the inflammation. (16) However, as of now, there is no definitive etiology for OLP, though some evidence suggests viral or bacterial infections, local trauma or irritation, systemic disorders, and even excessive alcohol and tobacco consumption are probable factors. (17)

Most of the current evidence supports the notion that OLP is related to immune dysregulation. However, the immune system is not only affected by drugs, systematic metabolic diseases, physical and mental stress, but also by microbiota and pathogens. The oral cavity is the beginning of the gastrointestinal system, and the gut-body connection and microbiome play important roles in maintaining health versus disease in the oral cavity and beyond. Therefore, OLP is likely a result of multiple factors which stress or dysregulate the immune system. By analyzing the gene expression profiles of cells in the oral cavity, we identified 953 genes that showed significantly different expression profiles in lichen planus patients compared to healthy controls. Among the differentially expressed genes, we identified 498 up-regulated and 455 down-regulated genes (p < 0.05, fold change > 2) between OLP patients and healthy controls **(Fig. 2A)**. To analyze the potential roles of the 953 differentially expressed genes, we used Ingenuity Pathway Analysis (IPA^®^) to identify the most likely molecular signaling mechanisms that could account for some of the symptomatic manifestations of OLP. Among the OLP patients, specifically enriched signaling pathways identified several genes coding for proteins involved in nicotine degradation (p= 0.003), cysteine biosynthesis and homocysteine degradation (p=0.015), as well pathways associated with increased activation of hepatocyte nuclear factor alpha (HNF4A) (p=0.015) and the modulation of its various downstream signaling molecules (**Fig 2B**). HNF4A is highly expressed in lymphatic tissue and in the salivary glands, which is consistent with the observation of an activated HNF4A gene network. Among the genes in this network, the largest difference in expression was observed in the sterol O-acyltransferase 2 gene (*SOAT2*) (increased 9-fold; p=0.005), a protein involved in cholesterol metabolism that has been implicated in various types of cancer including epithelial and liver cancer as well as hepatocellular carcinoma, melanoma, and acute myeloid leukemia. In addition to *SOAT2, CYP2C8* was upregulated (2-fold; p=0.03) and is an unspecific monoxygenase. Selectin P (*SELP*) was also upregulated (2-fold; p=0.004) and is a cell adhesion protein upregulated in response to bacterial infections, delayed hypersensitivity reaction, T-cell lymphoma, and systemic lupus erythematosus.

**Fig 2.**
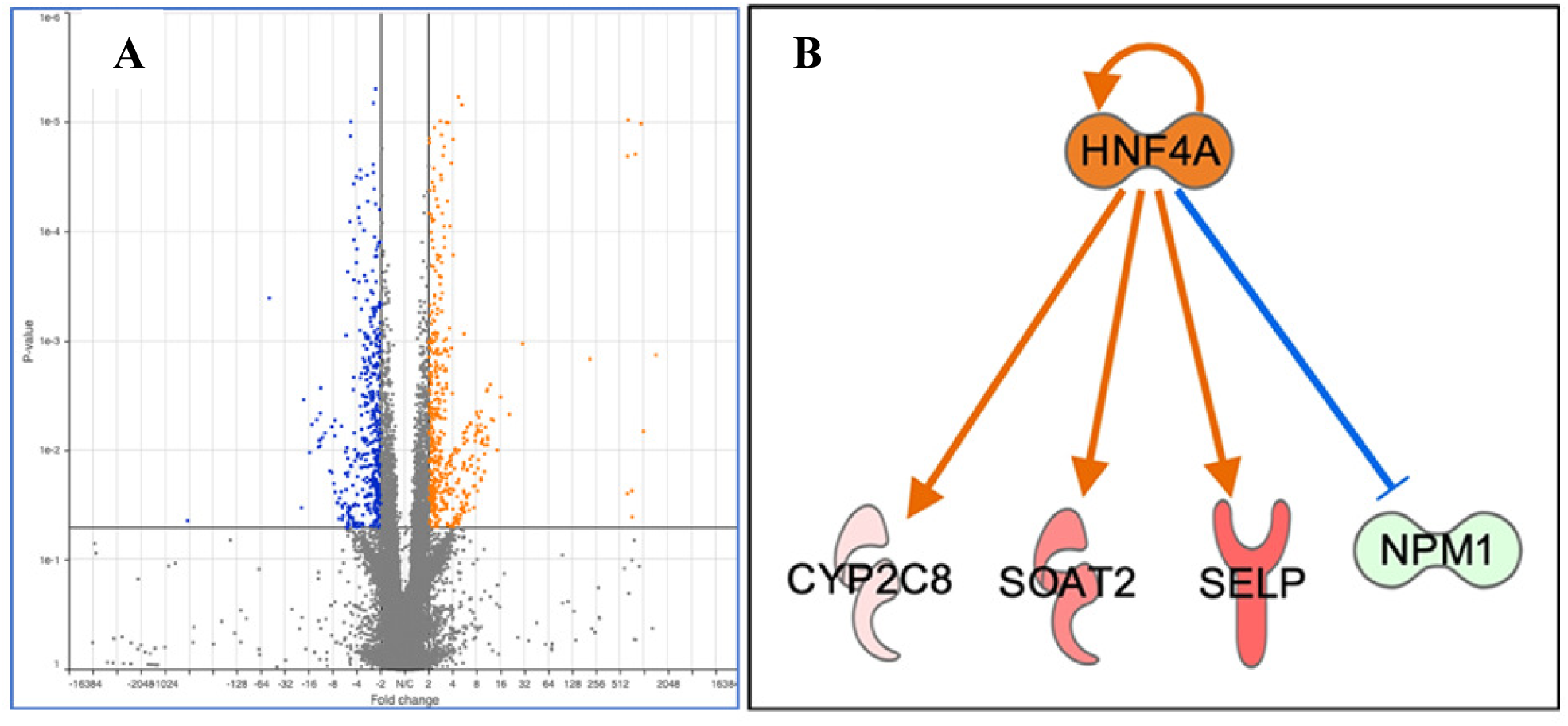
Differential expressed genes between OLP patients and healthy individuals. **A.** Volcano plot illustrating differentially expressed genes identified between OLP patients and healthy controls. Overall, we identified 953 differentially expressed genes of which 498 were upregulated and 455 were downregulated between OLP patients and controls. Further analysis of the differentially expressed genes identified hepatocyte nuclear factor 4 alpha (HNF4A) as a significant upstream regulator of the identified differentially expressed genes. **B.** The hepatocyte nuclear factor alpha (HNF4A) gene network is activated in OLP patients. Differential gene expression analysis indicated that the HNF4A gene network is activated in OLP patients. Comparing to healthy individuals, HNF4A and its down stream targets, CYO2C8, SOAT2 and SELP are upregulated in OLP samples, while the inhibitor of HNF4A, NPM1 is down-regulated in OLP compared to healthy individuals. Red: upregulation in OLP, Green: Down-regulated in OLP.

### Microbes associated with OLP

Considering that a typical oral cavity contains more than 500 different bacterial species and various viruses and yeast (18), the oral microbiome may play a significant and as-yet-undetermined role in OLP etiopathogenesis. Current next-generation sequencing (NGS) techniques provide powerful tools that can be used to identify the potential pathogens of OLP and facilitate new potential therapies. In particular, RNAseq can simultaneously identify microbial and host gene expression. As a proof of concept, we performed molecular analysis using RNAseq to compare OLP patients to healthy controls **(Table 1)** and we were able to identify putative candidates for OLP pathogenesis **(Fig 3)**. Of the identified microorganisms, 25 bacterial species were observed to be differentially expressed between OLP patients and healthy controls. A higher prevalence of *Corynebacterium matruchotii, Fusobacterium periodonticum, Streptococcus intermedius, Streptococcus oralis* and *Prevotella denticola* was found in patients with OLP as compared to controls (Fig 4). Clinically, these pathogens are significant, and in the context of OLP it has been established that patients with poor oral hygiene and plaque containing periodontopathogens respond poorly to conventional treatments for their OLP until their hygiene and plaque are adequately controlled. We also utilized PCR to confirm that *C. matruchotii, F. periodonticum, S. intermedius, S. oralis*, and *P. denticola* are dominant bacteria in OLP patients versus controls **(Fig 4)**. Finally, we also identified a higher abundance of viruses in OLP patients than in healthy controls. Three viral species were statistically significantly higher in OLP patients than in controls **(Fig 5)**, namely tick-borne encephalitis virus, bacillus virus SPO1, and a brochothrix bacteriophage virus.

**Fig 3.**
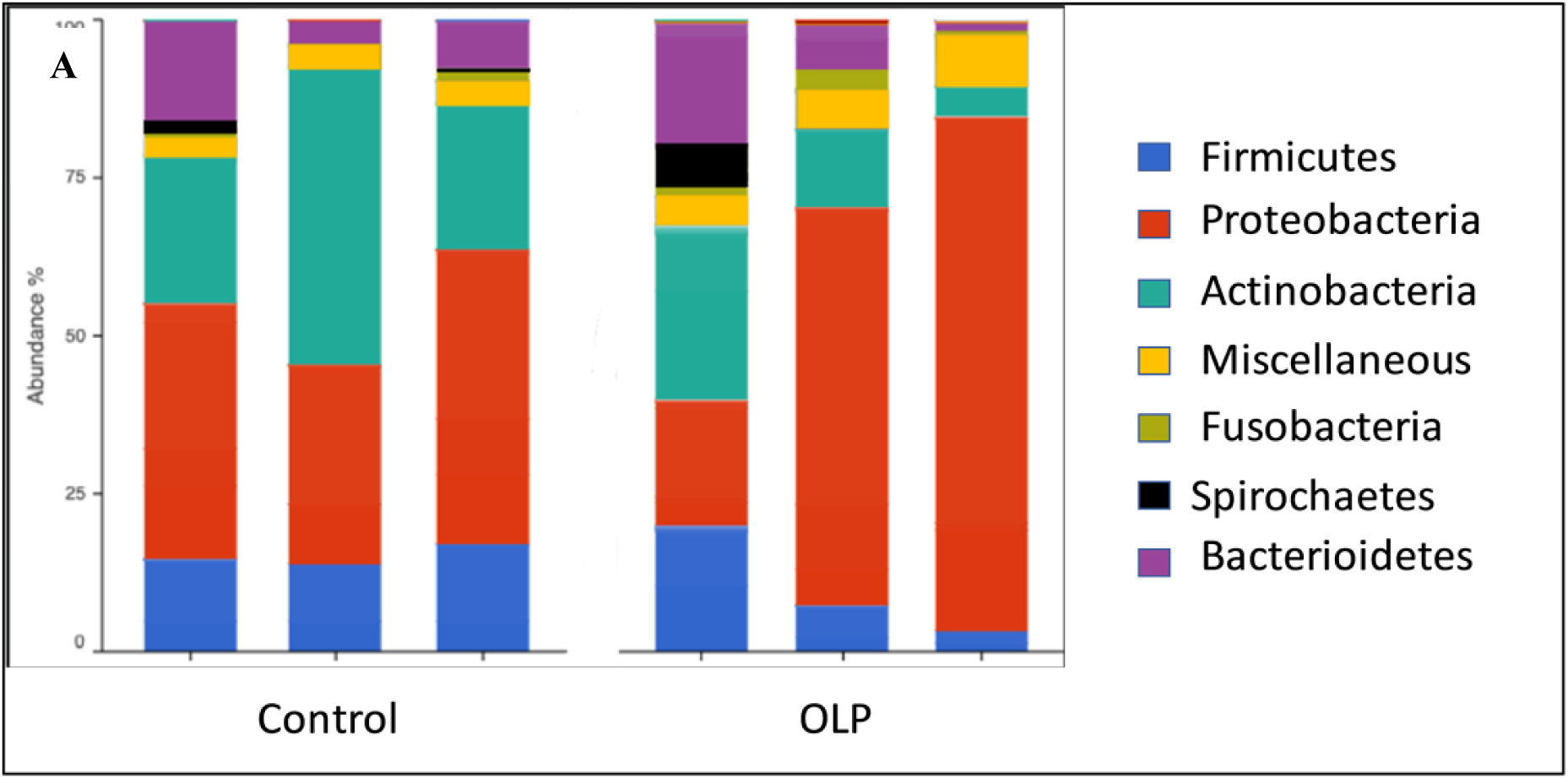
Relative abundance of identified bacterial and archaea species sequences in the saliva of patients with OLP and healthy controls. Overall an average, of 18 Million classified reads were recover from the RNA libraries and 2 million from the DNA, with no significant difference in microbial flora diversity. Among all samples, 1575 different bacterial species were identified. Of the identified microorganisms 25 bacterial species were observed to be differentially expressed between the OLP patients and healthy controls.

**Fig 4.**
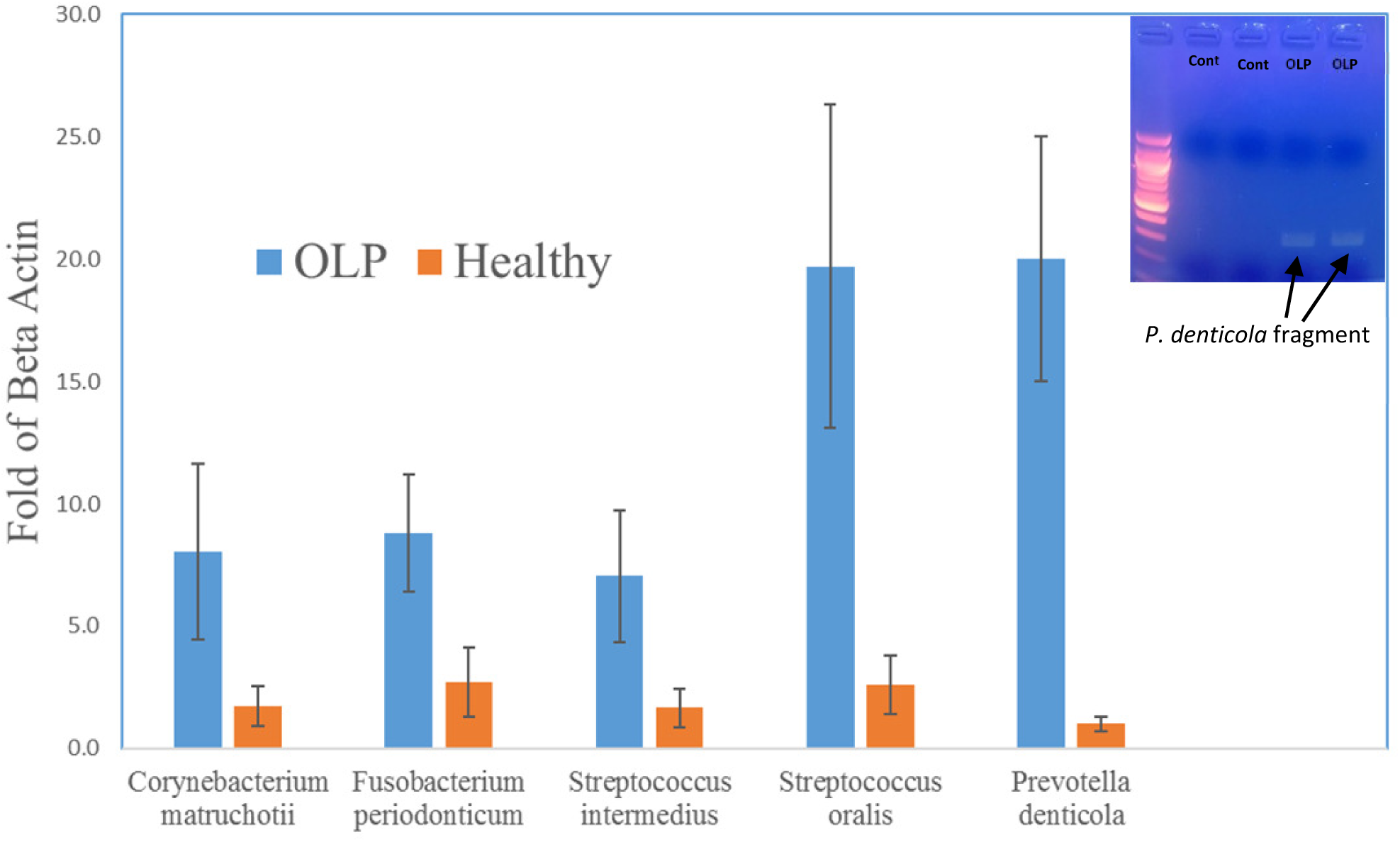
PCR confirmation of *Corynebacterium matruchotii, Fusobacterium periodonticum, Streptococcus intermedius, Streptococcus oralis* and *Prevotella denticola* are dominant bacteria in OLP patients (OLP: n= 10, Cont: n= 5). As a representative case, PCR primers designed to amplify a 316bp fragment of *Prevotella denticola* demonstrate the present of this bacteria in two OLP patients but not in matched healthy controls (Inset).

**Fig 5.**
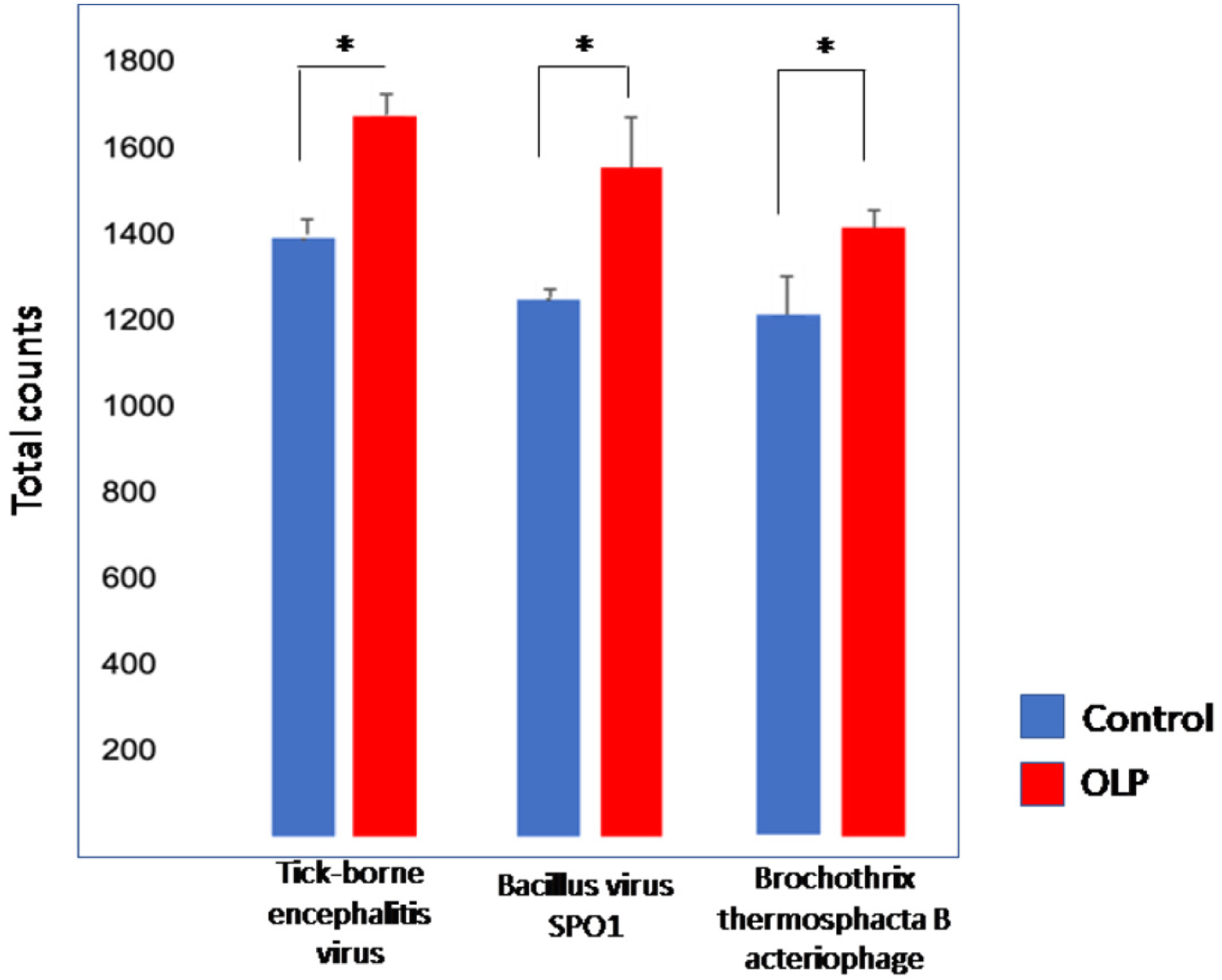
Three viral species are significantly different between OLP patients and healthy controls. P values are 0.0012, 0.0074, and 0.036 respectively from left to right, error bars represent s.e.m. Virus counts for the majority of species identified was higher in OLP patients when compared to that of controls.

## DISCUSSION

Since the causes of OLP have not been fully determined, there is no definitive cure for the condition. The first-line drugs in the treatment of OLP are topical corticosteroids due to their ability to modulate inflammation and immune responses by reducing the lymphocytic exudate and stabilizing the lysosomal membrane. (19) If topical steroids are not able to provide clinical relief and resolution or remission of painful erosive lesions, then systemic corticosteroids may be administered. (20, 21) Immune suppressants such as cyclosporine, a calcineurin inhibitor, are also used to reduce symptoms of inflammation and irritation.(20) Calcineurin is a protein phosphatase which is involved in the activation of transcription of IL-2 and stimulates the growth and differentiation of the T-cell response. By using calcineurin inhibitors, inflammation driven by T-cells, a suspected catalyst in the development of OLP, can be stopped. (19)

However, because knowledge of the underlying mechanism of OLP is lacking, current therapies are mainly prescribed based on their empirical results. In immunosuppressive therapy, topical calcineurin inhibitors (TCI) including tacrolimus, pimecrolimus, and cyclosporine are still controversial for use in this setting. Side effects include oral candidiasis, bad taste, nausea, dry mouth, sore throat, and swollen mouth but are considered minimal.(22) Topical retinoids such as tretinoin, isotretinoin, fenretinide, and tezarotene are generally less effective than topical corticosteroids and are more likely to cause adverse side effects.(23) When considering topical retinoids for treatment, the positive effects should be weighed against their rather significant side effects like cheilitis, elevation of serum liver enzymes and triglyceride levels, and teratogenicity. (19) Biologics and disease-modifying anti-rheumatic drugs (DMARDs) are also used in this setting, usually for cases that do not respond to steroid therapy.

It is critical to identify molecular pathways active in saliva and oral cavity cells of OLP patients and the related pathogens in order to improve OLP treatment and achieve resolution. Here, we found activation of the HNF4A gene network in oral cavity cells of OLP patients (Fig 6), and we further identified several periodontopathogens, including *Prevotella denticola*, which dominated in patients with OLP. Hepatocyte nuclear factor-4-alpha (HNF4A) is a member of the nuclear receptor superfamily of ligand-dependent transcription factors(24) and is the most abundant DNA-binding protein in the liver, where it regulates genes largely involved in the hepatic gluconeogenic program and lipid metabolism.(25) Besides the liver and pancreas (26), it is highly expressed in the kidney where it is involved with drug metabolism, (27) and also in the small intestine and colon where it is involved in inflammation. (28, 29) HNF4A is also known to affect inflammation and immune pathways in other immune-mediated conditions like Crohn’s disease (30) and inflammatory bowel syndrome.(28)

The HNF4A network could explain previously elusive associations of OLP with various diseases. Previously, chronic liver disease and hepatitis C have been found to be associated with OLP along with the presence of HLA-DR 6 (31) and HLA-A3. (32) Conditions like hypertension and diabetes mellitus also tend to have comorbidity with lichen planus.(20, 33) These associations may be due to disease effects on the HNF4A gene network in modulating the inflammatory and immune pathways of cells in the oral cavity. HNF4A network activation also explains the association of OLP or OLP-like (lichenoid) reactions with specific pharmacotherapies including beta blockers, nonsteroidal anti-inflammatory drugs, anti-malarials, diuretics, oral hypoglycemics, penicillamine, and oral retroviral medications. (20) These interesting associations of drug intake with OLP may be due to the systematic alteration of the HNF4A gene network and in the liver and kidney where it is involved in drug metabolism. (27)

We also identified a potential major pathogen, *P. denticola*, which could activate the HNF4A network in cells of the oral cavity. *P. denticola* may serve as a target for future therapies in this context, though further and larger studies will be needed to more reliably assess and validate this association. *Prevotella* species are found in humans as opportunistic pathogens. (34) More than twenty identified species of *Prevotella* are known to cause infection. (35) *P. denticola* has been isolated from the human mouth, where it is suspected to cause disease, (36) but its pathogenesis in OLP has remained elusive until this study. It has been reported that increased *Prevotella* abundance is associated with augmented T-helper type 17 (Th17)-mediated mucosal inflammation.(37) It has also been reported that *HNF4A* mutation is associated with Th17 cell-mediated inflammation of the gut mucosa. (38, 39) Taken together, our findings support the hypothesis that *P. denticola* abundance in the oral cavity can lead to activation of the HNF4A gene network in cells, resulting in Th17-mediated mucosal inflammation and diseases like OLP.

We also found greater viral abundance in OLP patients compared to healthy controls. Interestingly, OLP can have a clinical course similar to viral diseases affecting the oral cavity: specifically, the waxing and waning nature of the disease with bouts of active lesions and symptoms, followed by periods of quiescence or remission. Additionally, bacterial and viral interactions are key to many pathologic conditions, including those that involve the oral cavity. Certain bacteria have been shown to increase lytic replication of oral herpesvirus pathogens and thus polymicrobial interactions are likely key to disease pathogenesis. We have previously characterized a significant number of bacteriophage viruses in oral infectious diseases such as osteomyelitis and osteonecrosis, and such profiles were more abundant in diseased patients than in healthy controls.(40) Although the present work represents a preliminary small-scale study, we found a higher abundance of bacteriophages in OLP patients compared to controls. These findings taken together suggest that bacterial and viral interactions are important to oral disease processes that have inflammatory components, and OLP may be another such disease modulated by complex microbial and host interactions.

